# Vimentin promotes collective cell migration through collagen networks via increased matrix remodeling and spheroid fluidity

**DOI:** 10.1101/2024.06.17.599259

**Authors:** Minh Tri Ho Thanh, Arun Poudel, Shabeeb Ameen, Bobby Carroll, M. Wu, Pranav Soman, Tao Zhang, J.M. Schwarz, Alison E. Patteson

## Abstract

The intermediate filament (IF) protein vimentin is associated with many diseases with phenotypes of enhanced cellular migration and aggressive invasion through the extracellular matrix (ECM) of tissues, but vimentin’s role in in-vivo cell migration is still largely unclear. Vimentin is important for proper cellular adhesion and force generation, which are critical to cell migration; yet the vimentin cytoskeleton also hinders the ability of cells to squeeze through small pores in ECM, resisting migration. To identify the role of vimentin in collective cell migration, we generate spheroids of wide-type and vimentin-null mouse embryonic fibroblasts (mEFs) and embed them in a 3D collagen matrix. We find that loss of vimentin significantly impairs the ability of the spheroid to collectively expand through collagen networks and remodel the collagen network. Traction force analysis reveals that vimentin null spheroids exert less contractile force than their wild-type counterparts. In addition, spheroids made of mEFs with only vimentin unit length filaments (ULFs) exhibit similar behavior as vimentin-null spheroids, suggesting filamentous vimentin is required to promote 3D collective cell migration. We find the vimentin-mediated collective cell expansion is dependent on matrix metalloproteinase (MMP) degradation of the collagen matrix. Further, 3D vertex model simulation of spheroid and embedded ECM indicates that wild-type spheroids behave more fluid-like, enabling more active pulling and reconstructing the surrounding collagen network. Altogether, these results signify that VIF plays a critical role in enhancing migratory persistence in 3D matrix environments through MMP transportation and tissue fluidity.

## 1. Introduction

The collective motion of cells through tissue is a critical process underlying proper tissue maintenance, but dysregulated migration also contributes to diseases, such as fibrosis and cancer. In vivo cell migration is a complex, multi-step process that involves interactions between different cell types and their interactions with the extracellular matrix. Fibroblasts are the primary cells responsible for producing and remodeling the extracellular matrix (ECM) in homeostatic conditions and in response to injury [1–4]. They also are abundant in tumor tissue and promote the growth and invasion of cancer cells [5–7]. For efficient cell migration, fibroblasts secrete matrix metalloproteinases (MMPs) that degrade the ECM structure and create more space for other cells to move as well as generate large contractile forces that align collagen fibers, which promote guided migration and wound repair.

Vimentin is a type III intermediate filament protein expressed in cells of mesenchymal origin, including fibroblasts. Vimentin is important for wound healing, but it is also highly expressed in metastatic cancers and tumors, correlating with poor patience prognosis [8–10]. Vimentin is also a widely used marker of the epithelial-to-mesenchymal, in which cohesive epithelial cells lose their cell-cell contacts and transition to a polarized, motile mesenchymal phenotype [11]. Vimentin is perhaps best known for its cytoskeletal role as a mechanical scaffold that gives strength to the cell [12–14], but it is also known for its functional diversity and signaling roles associated with cell adhesion, organelle transport, cell polarization, and nuclear positioning. Despite these observations, the role of vimentin in cell migration through ECM environments remains poorly understood.

In many ways, the vimentin phenotype has been elusive. In 1994, the vimentin-null mouse was reported to develop and reproduce without an obvious phenotype [15], though by 2000, it was found that the vimentin-null mouse had impaired wound healing abilities [16]. Since then, there have been at least 39 phenotypes revealed by the vimentin-null mouse linked to a variety of functions, ranging from tissue remodeling, vascular functions, fat accumulation, tumorigenesis, and viral and bacterial infections [17]. Prior studies have hinted at mechanistic insight for vimentin’s role in collective cell migration relevant to in vivo conditions. For instance, vimentin promotes polarization and collective cell closure of wounds in 2D scratch wound assays [18, 19] and also increases collagen network contraction, at least for relatively low cell concentrations with few cell-cell contacts [20, 21]. Recent studies, however, paint a more nuanced picture because at the single cell level, vimentin decreases 3D cell migration speeds [22] and can also reduce single cell contractility [23–25]. Further, A recent study by Ostrowska-Podhorodecka et al [26] also revealed a new role of vimentin in transport of the membrane type 1 (MT1)-MMP, whose role in degrading collagen may be important for making room for collective cell migration through collagen networks. Taken together, the main mechanism behind collective cell migration by fibroblasts through a surrounding collagen remains unclear.

In this manuscript, we use a three-dimensional model of cell spheroids embedded in collagen I matrices to probe the role of vimentin in collective cell migration. Collagen I is the main component of the ECM in connective tissues and solid tumor [27, 28], and gels formed by collagen I are a common model system for the tissue microenvironment [29, 30]. Spheroids comprised of fibroblasts mimic physiological conditions in which fibroblasts detached from connective tissues, during inflammation, wound healing, and cancer cell migration [31]. To investigate the effects of vimentin on collective cell migration in 3D ECM environments, we developed and examined spheroids of wild-type and vimentin-null mouse embryonic fibroblasts embedded in collagen I matrixes with and without co-culturing with breast cancer cells, MDA-MB-231. We find that vimentin significantly promotes the collective migration through collagen for both fibroblast spheroids and spheroids of mixed populations of fibroblast and cancer cell spheroids. To understand these effects, we further examine the role of vimentin network structure, spheroid contractility, and collagen degradation activity on vimentin-mediated collective cell migration. Our results show an intact vimentin cytoskeleton is a significant contributor to collective cell invasion in collagen network and suggest new mechanisms by which vimentin mediates collective cell spreading and motility in real tissue.

## 2. Results

### Vimentin enhances collective cell migration through collagen networks

To investigate the role of vimentin in 3D collective cell migration, we developed fibroblast spheroids comprised of either wild-type or vimentin-null mEF. Spheroids were generated by culturing suspended cells in non-adhesive wells for 48 hours (Methods) (Fig. 1), creating spheroids 100-200 μm in diameter. Spheroids were then collected and embedded in collagen I matrixes (collagen concentration = 1.5 mg/mL).

**Figure 1.**
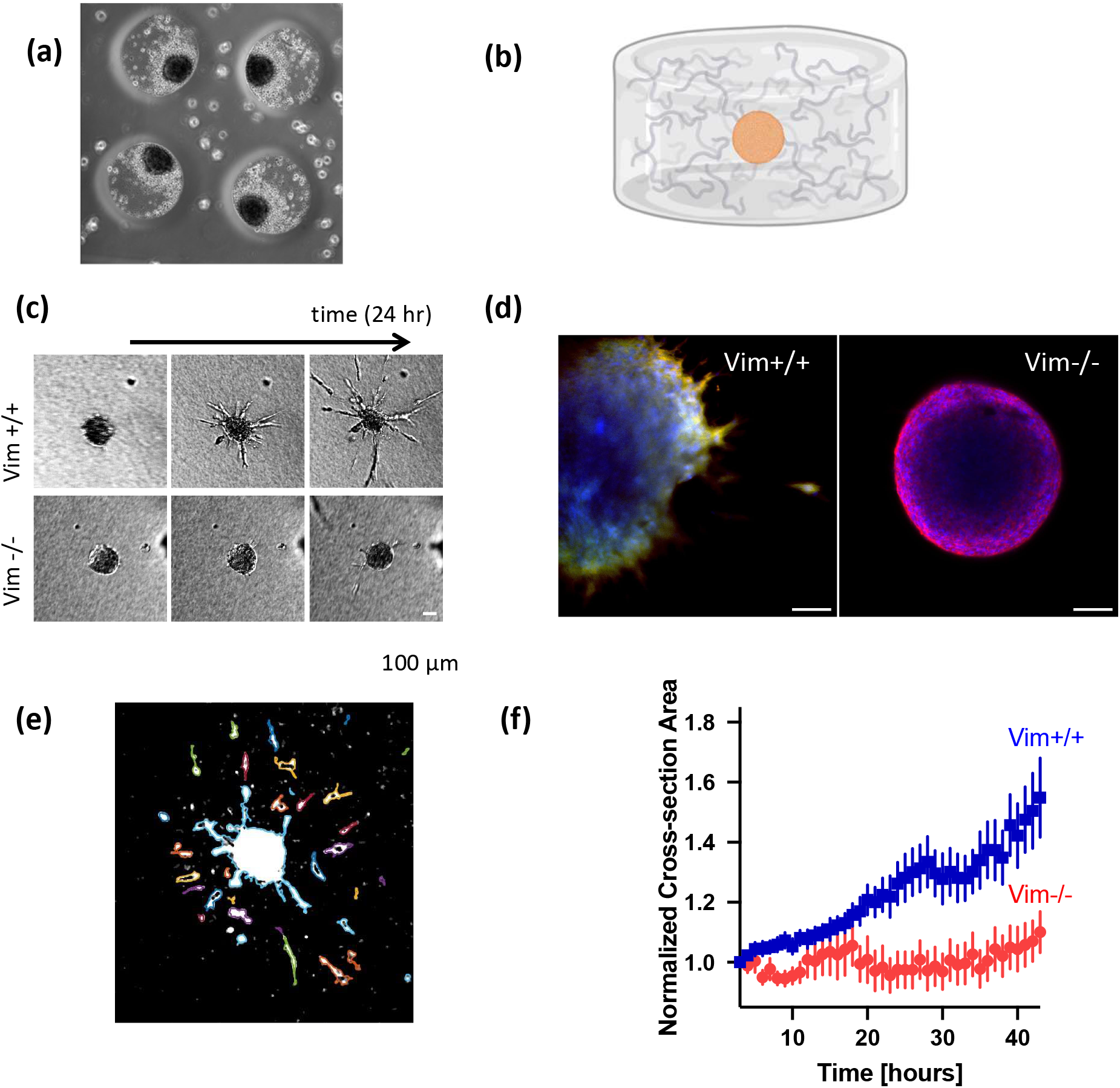
A) Spheroids formation in fabricated microwell (r = 200 um). B) Schematic of embedded spheroid in collagen gel (1.5mg/ml). C) Z-projected confocal image of Vim+/+ and Vim-/- spheroid in collagen gel (Blue:Nucleus, Green: Vimentin, Red: Actin), D) Images of Vim+/+ and Vim-/- spheroid in collagen gel over 24 hours. E) Cross-sectional area analysis of the spheroid in MATLAB. Area of the spheroid only count the core and visibly attached protrusion. E) Normalized cross-sectional area comparison between Vim+/+ and Vim-/- spheroids over 48 hours. Vim+/+ spheroids expand into collagen gel 60% more than Vim-/- spheroid. (n ≳ 4 independent experiments, N≳ 24 spheroids)

Representative time-lapse images of the spheroid invasion behavior are shown in Figure 1D. Wild-type mEF spheroids display characteristic features of collective cell expansion in ECM network, in particular detachment of individual cells into the network as well as collective streams of cells radiating out from the core of the spheroid. The vimentin-null spheroids display very different behavior: cell migration into the surrounding network is strongly diminished. At 8 hours, some cell protrusions can be seen invading the network, those these protrusions are dynamic and many collapse back toward the spheroid core (SI Movies). While markedly less compared to wild-type mEF, a few vimentin-null cells can be seen to form stable protrusions into the network or detach by 48 hours.

To quantify the expansion, we identified and tracked the boundary of the cell spheroid using a custom image analysis thresholding code (Fig. 1E, Methods). We measured the mean projected area of spheroids, which includes the core of the spheroid and connected cell protrusions but not detached cells over time (Methods). Figure 1F shows the mean projected area of the spheroids normalized by their initial projected area (Methods). Vimentin increased the projected area of the spheroid, increasing by 60% compared to the spheroids of vimentin-null cells (p < 0.01). Overall, the results in Fig. 1 indicate that vimentin regulates 3D cell migration through collagen networks, promoting the expansion of fibroblast spheroids through an ECM network.

### Vimentin increases collagen network remodeling and spheroid contractility

Next, we quantified remodeling of the collagen network using reflective confocal imaging and traction-force microscopy-based techniques, as shown Fig. 2 and 3. Confocal reflectance microscopy uses back-scattered light to form an image and is commonly used for imaging collagen networks simultaneously with fluorescence from cells [32]. Figure 3 shows representative images of the spheroids after 48 hours in the collagen gel. The wild-type spheroids show a collagen pulling behavior with fibers radiating out from the spheroid, consistent with the remodeling of collagen associated with spheroid expansion through ECM networks [33–35]. In contrast, no radial fibers are observed in the vimentin-null spheroid. To analyze the collagen remodeling, we computed the compaction of the collagen network around the spheroids (Methods), which showed an increase in collagen compaction in the vimentin-containing spheroids by 50% (p <0.05). We also quantified the alignment of the collagen fibrils around the spheroids (Methods) and found an increase in collagen alignment 2 times in wild-type compared to vimentin-null spheroids (p<0.0001).

**Figure 2.**
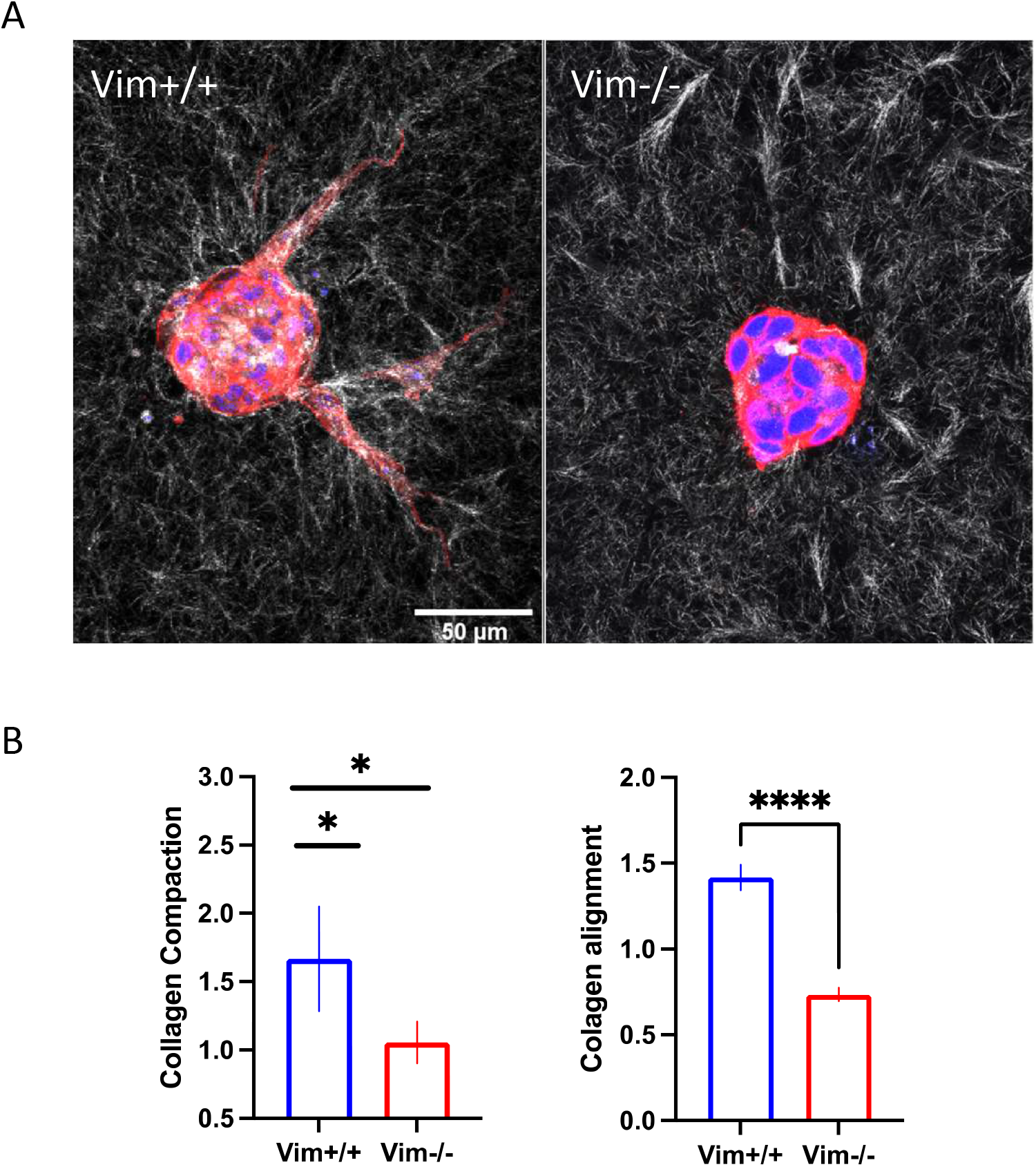
A. Confocal reflectance image of Vim+/+ and Vim-/- spheroid in collagen gel (Blue: DAPI, Red: Actin, White: Collagen) B. Collagen compaction and collagen alignment analysis of Vim+/+ and Vim-/- spheroids. Vim+/+ spheroids significantly compact and align the network compared to Vim-/- spheroids. (n ≳ 2, N ≳ 24)

**Figure 3.**
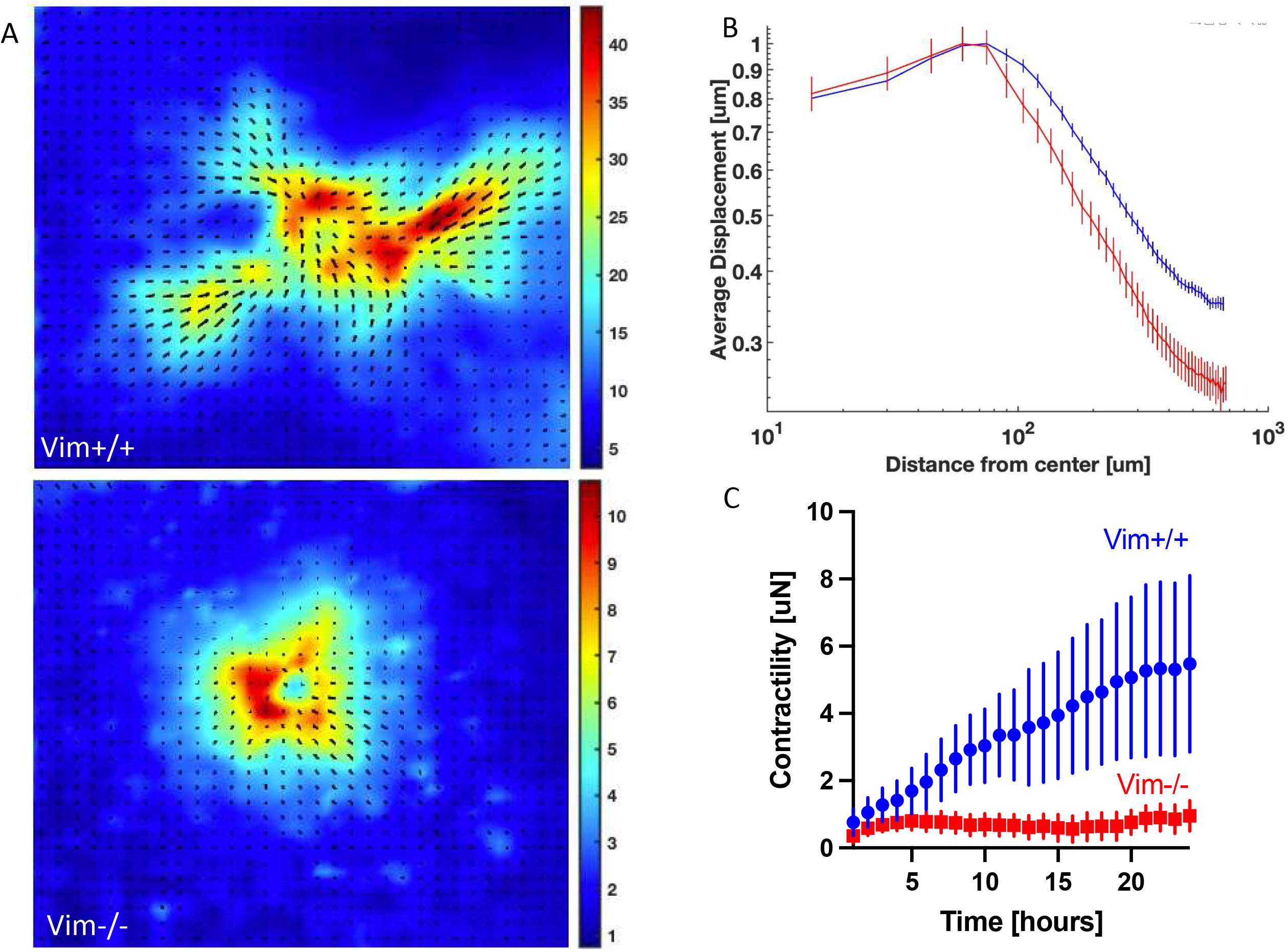
A. Aggregated displacement field and magnitude of a Vim+/+ and Vim-/- spheroid. B. Aggregated displacement as a function of distance away from the spheroid center. C. Contractility of Vim+/+ and Vim-/- spheroids over 48 hours. Vim+/+ spheroids contracts collagen gel more and further than Vim-/-. (n ≳ 3, N ≳ 13)

Traction-force microscopy-based experiments were conducted by embedding the collagen networks with 2 μm fluorescence beads and performing time-lapse imaging of the collagen networks around the spheroids over time (SI Videos). The videos show fluctuations of the tracer beads around the spheroids but a systematic net motion of the beads toward the spheroids over many hours, indicative of the generation of contractile pulling forces of the spheroid on the ECM network. To compare the wild-type and vimentin-null spheroids, we computed a time-averaged displacement field, as shown in Fig. 3B. The time-averaged displacement fields show a net inward contraction of the collagen toward the spheroids and the magnitudes of collagen displacement are greater for the wild-type compared to vimentin-null spheroids. An interesting feature of the wild-type fibroblast spheroids is nonuniform regions of increased contraction that correlate with regions of outward invading strands of cells. The spatial differences in collage displacements between the two spheroid types are highlighted by radially averaged displacement as a function of r from the center of the spheroid (Fig. 3C).

To connect the collagen deformation fields to forces generated by the spheroids, we use the finite element approach described in Mark et al [36] (Methods,SI). In this model, the spheroid is considered as a uniformly contractile sphere in a fiber-based network with mechanical properties and pore size parameters determined by experimental data. By fitting the experimentally measured collagen displacement data to the finite element results, we can extract an effective contractility that the spheroid exerts on the collagen network as a function of time. With this approach, we find that the contractility of the wild-type spheroids is significantly greater than the vimentin-null cell spheroids, as shown in Fig. 3C. In particular, the contractile forces of the wild-type cell spheroids is approximately 6 μN compared to 1 μN for the vimentin-null spheroids. These data indicate that vimentin increases the contractility of collective cell spheroids in a soft 3D collagen network, consistent with the increased migration of the wild-type cells seen in Fig. 1.

### Vimentin-mediated collective cell migration depends on physical vimentin networks

Our results thus far demonstrate vimentin plays an important role in collective migration of fibroblast cells though collagen networks. To investigate the whether the physical structure of vimentin networks is essential for the observed collective cell migration, we next studied the ‘unit length filament’ (ULF) vimentin mutant Y117L. The vimentin unit-length filament, consisting of approximately 8 vimentin tetramers and 60 nm in length, is considered the smallest stable vimentin structure inside the cell and is the basic repeating unit in vimentin polymers [37]. These ULF mutants are prepared by expressing ULF vimentin in vimentin-null cells and thus retain vimentin’s signaling interactions within the cell, but its assembly into long filaments is blocked at the ULF stage [38]. Figure 4A shows that the knockout cells expressing ULF vimentin segments did not rescue the expansion behavior of wild-type cells. These ULF cells migrated poorly into the matrix and generated low-contractility, similar to the vimentin-null spheroids. The data in Figure 4C and 4D suggests that intact vimentin networks are required for collective cell expansion in through collagen networks.

**Figure 4.**
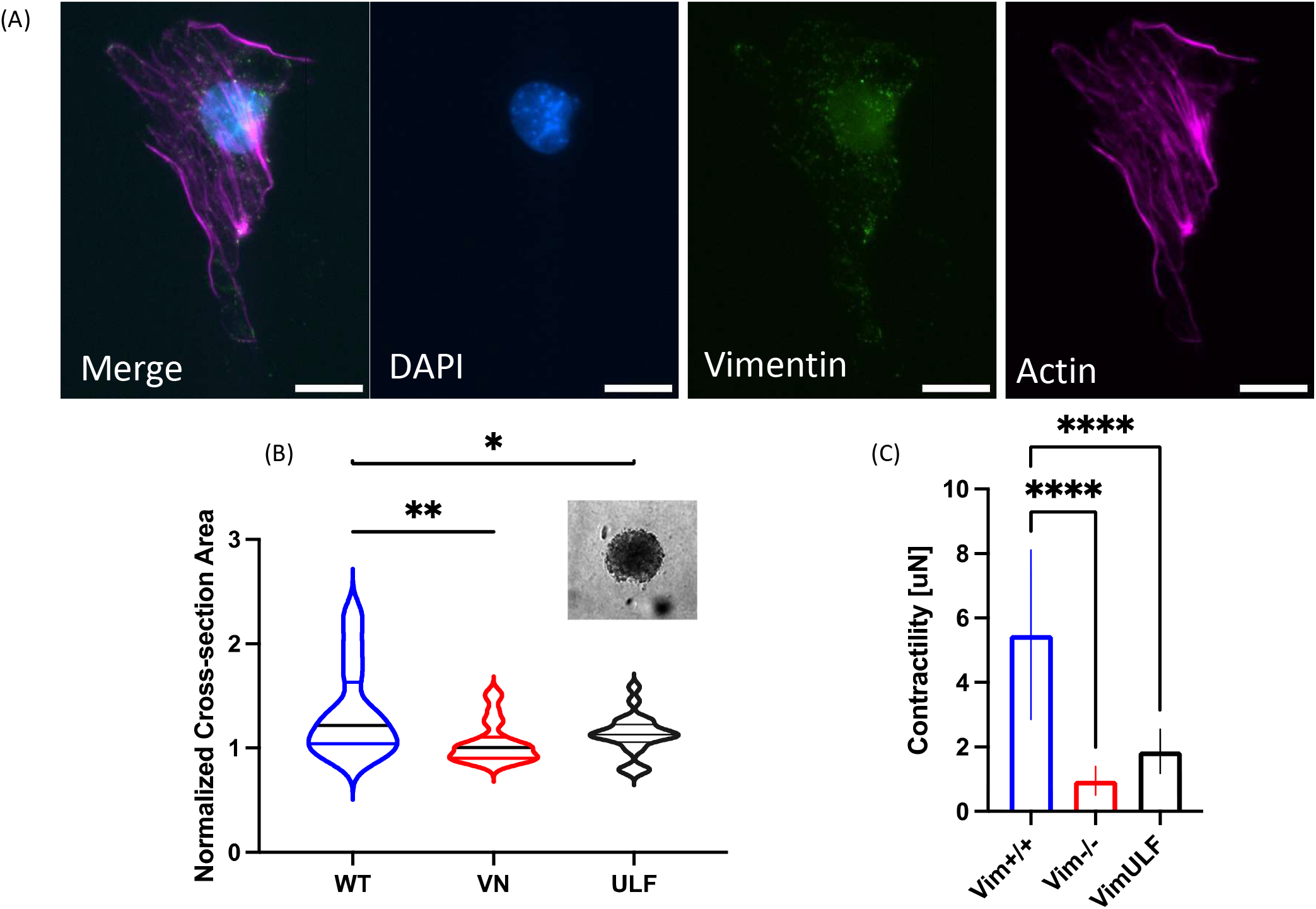
A) Fluorescent image of Vimentin-ULF fibroblasts (Blue: DAPI, Green: Vimentin, Magenta: Actin. B) Image of a spheroid made of VimULF fibroblasts embedded in Collagen gel (1.5mg/ml) at 48 hours. C) Normalized Cell Area and D) Contractility comparison between Vim+/+, Vim-/- and VimULF spheroids. VimULF cells exhibit similar phenotype to Vim-/- spheroids. They do not demonstrate high invasiveness and high contractility. (n ≳ 3, N≳ 22 for cell area expm, n ≳ 2, N ≳ 9 for contractility experiment)

### Vimentin-mediated migration is MMP-dependent

Prior studies highlight that the migration rates of cells through collagen strongly depend on matrix metalloproteinases (MMPs), which degrade ECM fibers and create space for migration. To determine the role of MMP in the spheroid expansion, we treated the embedded spheroids with 25 µm of GM6001 broad spectrum MMP inhibitor (Methods). The MMP inhibitor strongly suppressed expansion of wild-type cells into the collagen network and decreased spheroid contractility. It also limited the protrusive cellular structures seen in the vimentin-null cells.

Based on these results, we hypothesized that the decreased migration of vimentin-null cells could come from alterations in collagen degradation. Further, the vimentin-null cells may be capable of migrating into the collagen network if the confinement from the matrix was alleviated. To test this picture, we next performed experiments with laser-ablation to generate micro-tracks in the collagen network (Fig. 5). As shown in Fig. 5, the wild-type cells move out into the microtracks but also make their own additional tracks invading into the network. In contrast, the vimentin-null cells did not make their own tracks but did move out into the ablated collagen micro-tracks. This data suggests that the confinement of the collagen matrix here is a major barrier for the vimentin-null cells’ motion and that these cells are able to move in collagen when that barrier is removed.

**Figure 5.**
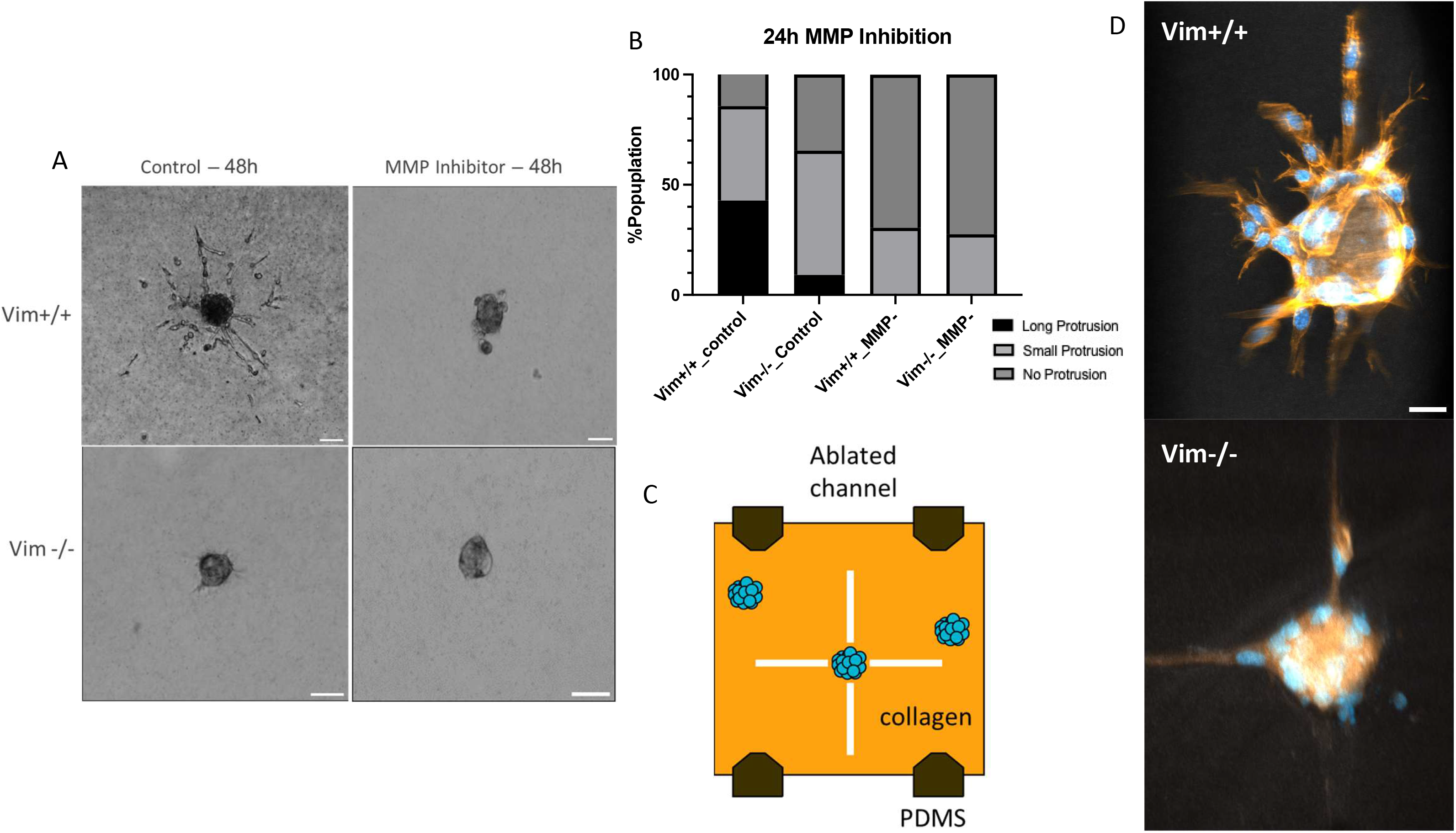
A) Image of Vim+/+ and Vim-/- spheroids at 48hours with and without GM6001 broad spectrum MMP inhibitor. B) Quantitative analysis of spheroids population that show protrusion. Long protrusion indicates strand invasion that is longer than 50um (approximately half of the spheroid radius). Small protrusion indicates that invasive morphology can be observed but the length of the strand is smaller than the radius of the spheroid. MMP inhibited spheroids are no longer capable of high invasion regardless of cell line. C) Schematic of laser ablated channels in collagen gel. D) Fluorescent image of the Vim+/+ and Vim-/- spheroid in ablated collagen gel. Vim-/- spheroids are capable of invasion into the ablated channels. E) Schematic showing relationship between Vimentin and MT1-MMP.

### 3D segmentation shows vimentin increases nuclear polarization in collagen-embedded cell spheroids

Our results thus far indicate that filamentous vimentin enhances fibroblast collective expansion and spheroid contractility, and these features are dependent on MMP-degradation of the collagen network. Degradation of the collagen network likely makes space for cells and cellular protrusion to expand into the collagen, allowing cells to better grip and pull-on collagen fibers to aid migration. However, another increasingly recognized aspect of collective cell migration is related to the fluidization of the collective cell aggregate, spheroids, or tumors [39–43]. In this context, the degree of fluidity of a collective cell aggregate refers to the ability of cells to rearrange and diffuse among their neighbors in the aggregate, allowing for motion and mechanical features akin to fluid flow. There has been much recent work on understanding this rigidity transition of different tissues, in other words, the transition from a stable solid-like structure to a fluid-like tissue, which is relevant to the collective motion of cellular aggregates in vivo. One finding that has been well described is that the distribution of cell shapes in epithelial-like tissues can predict where the aggregate is in relation to the rigidity transition, e.g. whether the aggregate will behave as a solid- or fluid-like material. More elongated cell shapes, based on a cell-shape index, are associated with more fluid-like behavior. Recent work indicates a similar transition occurs in 3D spheroid aggregates [41, 44], and that there is a close relation between cell shape and nuclear shape (which is experimentally much easier to measure in 3D spheroidal systems) [45, 46].

Motivated by these findings, we measured the distribution of 3D nuclear shapes in our wild-type and vimentin-null spheroids, which is technically more feasible than measuring 3D cell shapes in a spheroid. We considered two time points, 6 hours and 48 hours past embedding the spheroids into the collagen network to compare nuclear shapes distributions pre- and post-invasion into the network (Fig. 6). Nuclear shapes were assessed by running TWANG Segmentation code [47] on confocal images of nuclei from embedded spheroid. We extracted the major and minor axis of the nuclei data to calculate nucleus aspect ratio. (Methods). As shown in Fig. 6B, we found that the nuclei in the wild-type spheroid were significantly more elongated compared to the vimentin-null spheroid in both time points. The wild-type spheroids, in particular, exhibit more elongated nuclei, which tend to localize around the perimeter of spheroid, consistent with prior studies [41, 44, 48]. The wild-type distribution of nuclear shapes thus exhibits a maximum around 1.2, for the cells in the center of the spheroid, and a long tail at high nuclear elongation shapes for the boundary cells. At 48 hours, the vim+/+ distribution started to shift to the right, more nuclei can be observed with aspect ratio of 1.8 to 2.3 with some highly elongated nuclei reaching AR of 2.8.

**Figure 6.**
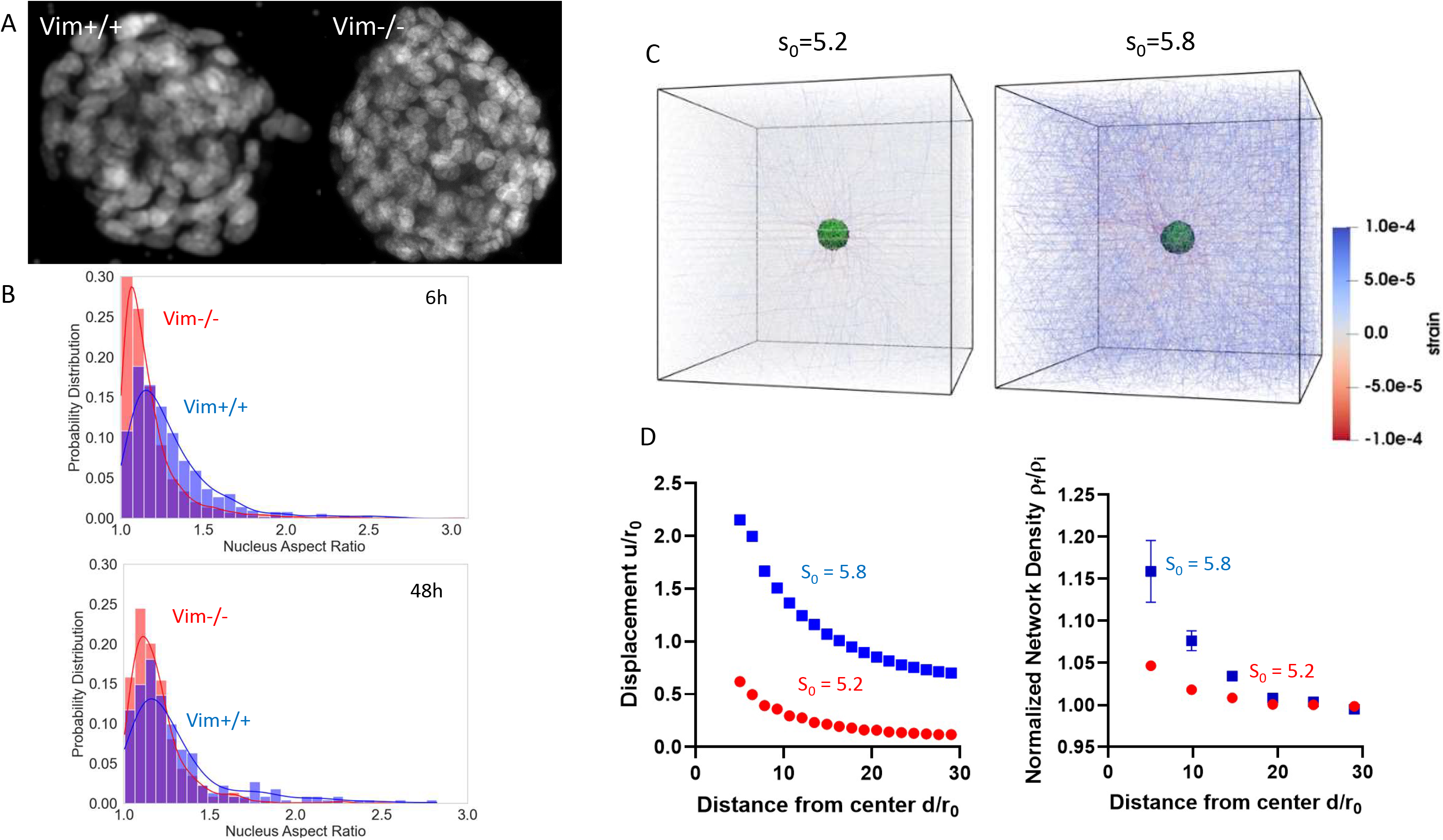
A. Segmentation image of Vim+/+ and Vim-/- nuclei inside embedded spheroid. B. Nucleus Aspect Ratio of Vim+/+ and Vim-/- spheroid at 6 hours and 48 hours. Vim+/+ nuclei becomes more elongated over time compared to Vim-/- nuclei. C. 3D spherical vertex model of a fluid-like (s_0_ = 5.8) and solid-like (s_0_ = 5.2) spheroids in a cross-linked network of fiber. S_0_ is the represents the dimensionless shape index, defined as 𝑠_0_ = 𝐴/𝑉^2/3^ where A is the cell surface area and V is the cell volume. Larger s_0_ indicates elongated cell shapes. D. Simulated normalized displacement and density of the network as a function of distance away from the center of fluid-like and solid-like spheroid. Fluid-like spheroids cause more network displacement and densify the network more compared to solid-like spheroid.

Taken together, these data point to an additional effect of vimentin on the collective cell expansion, namely vimentin increases nuclear elongation, a characteristic feature of increasing tissue fluidity. This connection exists at a pre-invasion stage (6 hours), either independently or in parallel with degradation of the surrounding collagen that proceeds invasion.

### Increased tissue fluidity enhances collagen remodeling

To establish an interpretive framework for our results (Fig. 6C), we invoked a novel 3D computational model of a cell spheroid embedded in a collagen network [49] and compared the predictions of the model to our experimental results. As shown in Fig. 6C, the spheroid is comprised of a 3D spherical vertex model, in which cells are represented as deformable polyhedrons that share faces with neighboring cells. Cells on the periphery of the spheroid are connected to a collagen matrix, which is well represented as a cross-linked network of fibers, and cells in the spheroid actively pull on the fibers connected to them. While prior literature work has developed models of contractile spheroids in collagen [36, 50, 51], the 3D vertex spheroid model allows connections between individual 3D cell shapes and the remodeling of the collagen network around it. Vertex modeling of biological tissues has been a useful approach in capturing the dynamics of 2D cell systems and 3D spheroids in fluid (not embedded in collagen). An advantage of this approach is its potential to generate quantitative predictions on cell mobility and tissue fluidity based on information of cell shape alone, e.g. elongated cell shapes reflect higher cell motility whereas less elongated, more spherical cell shapes indicate an arrested, less motile cell state [41, 49, 52, 53]. In the 3D model, the cell shape is assessed through s, a dimensionless shape index that depends on the cells target surface area 𝐴 defined as 𝑠 = 𝐴/𝑉^2/3^, where 𝑉 is the volume of the cell. Larger s indicates more elongated cell shapes. A current limitation of the model is the lack of explicit collagen-degradation and cell-break; however, this minimal model is a useful starting point to examine how changes in cell shape (as indicated in Fig. 6B) impact collagen remodeling.

Figure 6C shows computational pictures of spheroids embedded in collagen networks, for spheroids with target cell shape indexes of 5.2 and 5.8. The simulation parameters used here are the same as in Tao Zhang et al. [49] but with a fiber edge occupation probability p=0.85. The data is shown for time point sufficient large enough to allow for both the cells and fibers to deform under the cell pulling forces to rearrange and reach a new equilibrium that differs from the initial state (SI). The pictures show that collagen remodeling and network strain significantly increases for the high shape index s=5.8 compared to 5.2. To quantify this effect, we measured the collagen concentration, the collagen displacement, and collagen strain, as a function of the radial distance away from the center of spheroid. All three of these metrics – which quantify different aspects of collagen remodeling – increases for spheroid with increased cell elongation, as shown in Fig. 6B. These data indicate that more polarized cells, reflective of a more fluid-like spheroid, pull on and rearrange surrounding collagen to a greater extent than non-polarized cells, reflective of a more solid-like spheroid. This effect is the result of increased cell dynamics and cellular rearrangement that emerge over time in the fluid-like spheroid, allowing cells to increase their frequency of contact and pulling on the collagen surrounding the spheroid [40].

This computational model seems to capture main features of the experiments data and connect cell shape with collagen remodeling. Namely, we find more polarized cells - as indicated by the elongated nuclear shapes in wild-type cells – exhibit more collagen remodeling in the spheroid-collagen model presented here (Fig. 6D). These data give rise to the intriguing hypothesis that vimentin increases collective cell migration by (at least in part) polarizing cells and increasing tissue fluidity, which supports enhanced collagen remodeling at the collective cell level. Because the compaction and cell-induced stress of a collagen network feeds back into cell migration [54], this effect further promotes the expansion of the cells into the collagen network, in addition to polarization and fluidization of the cells themselves. More broadly, our experimental and computational data supports the inference that a more fluidized spheroid (as indicated by elongated nuclear shape) more strongly rearranges a surrounding collagen network, which to the best of our knowledge has not been previously demonstrated in this detail and is an unrecognized aspect of spheroid expansion in collagen networks.

### Vimentin intermediate filaments aid collective escape of fibroblast-cancer co-cultures

The experiments so far indicate a role of vimentin in enhancing collective migration of fibroblasts in collagen gels, which may have important physiological consequences. For example, fiibroblasts play an important role in growth and expansion of tumors into the surrounding soft matrix environments [5–7, 55]. To test whether vimentin impacts fibroblast function in cancer cell migration, we conducted experiments with fibroblast-cancer cell co-cultures. Specifically, we generated spheroids comprised of both mEF and the breast cancer cells MDA-MB-231 (SI, Methods), at a ratio of 2:1 consistent with prior co-cultures models [56, 57]. We found that the presence of vimentin in the fibroblasts of the co-cultures had a significant impact on the spheroid expansion. Figure 7 shows representative images of spheroids comprised of (i) MDA-MD-231 cells alone, (ii) wild-type mEF and MDA-MB-231 cells combined and (iii), vimentin-null mEF and MDA-MB-231 cells. Notably, the wild-type mEF and MDA-MB-231 cells showed increased expansion compared to the MDA-MB-231, pointing to the role of fibroblasts in aiding cancer cell migration. In a striking contrast, we found that the co-cultures of vimentin-null mEF and MDA-MB-231 expansion was almost entirely suppressed. The data in Figure 7F indicates that vimentin has a strong impact in the function role of fibroblasts in co-culture cancer cell migration, which could have important therapeutic applications for treating cancer by targeting vimentin.

**Figure 7.**
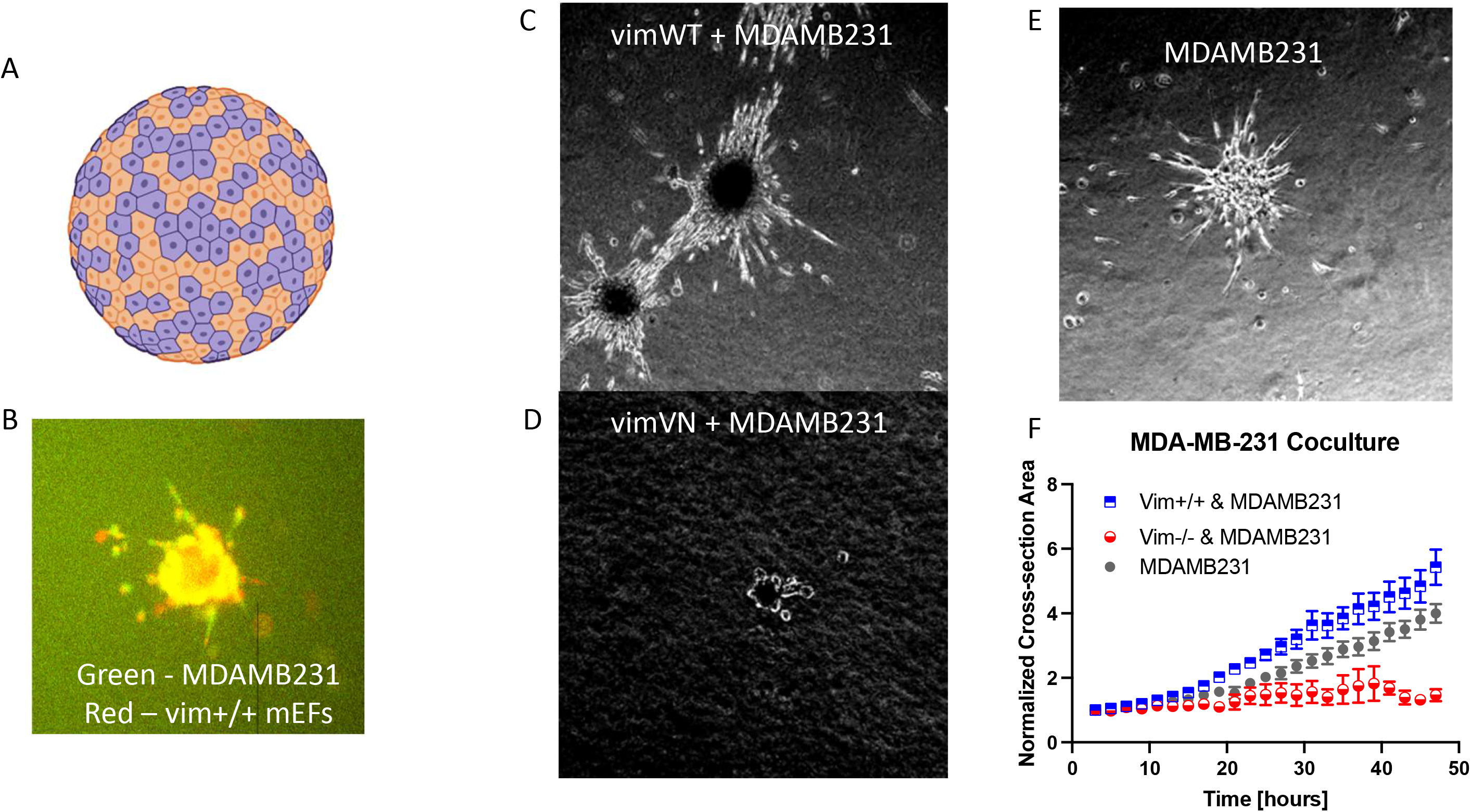
A) Schematic of coculture spheroids, B) Membrane-live-stained of coculture MDA-MB-231 and mEFs spheroids (Green: MDA-MB-231, Red: mEFs). C) Morphology of coculture spheroid at 48h of Vim+/+ mEFs and MDAMB231, D) Coculture spheroid morphology at 48h of Vim-/- mEFs and MDAMB231. E) Morphology of MDAMB231 spheroid. Normalized area comparison of the coculture spheroids and monoculture cancer spheroids over 48h. Coculture with Vim+/+ mEFs enhances the invasiveness of the cancerous spheroid. (n = 2, N ≳ 9)

**Figure 8.**
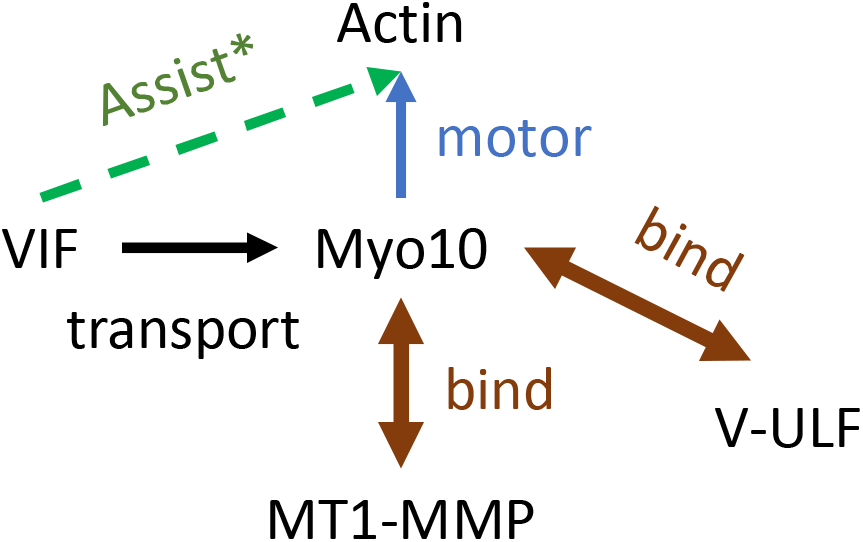
Relationship schematic between Vimentin and MMP. MT1-MMP binds to myosin 10 which is an actin-based motor. Myosin 10 is transported to the cell boundary via vimentin intermediate filament network. Hence a robust vimentin intermediate filament network allows effective transportation of MT1-MMP to the cell membrane, which results in better degradation of collagen in vimentin wild-type spheroids.

## 3. Discussion and Conclusion

Our results demonstrate that the vimentin cytoskeleton plays an important role in driving collective cell migration through ECM networks, as indicated by spheroids comprised of both fibroblasts alone and fibroblast-cancer cells co-cultures. This increased cell migration correlates with increased spheroid contractility, protease-mediated collagen degradation, and spheroid fluidity. Further, disrupting the assembly of filamentous vimentin networks suppresses collective cell migration through collagen, indicating the presence of filamentous vimentin is necessary for collective fibroblast migration through ECM networks.

The observed increase in motility in Vim+/+ spheroid is not surprising, given that repeated experiments showed a higher collective migration of cells expressing vimentin in 2D wound healing assay [58]. Eckes et al. have shown that fibroblasts derived from transgenic KO mice deficient for vimentin showed delayed migration of fibroblasts into the wound site [16]. Cheng et al. have demonstrated that loss of vimentin leads to a severe deficiency in fibroblast growth and abolished collagen accumulation in the Vim-/- wounds [59]. Walker et al. showed that vimentin knockdown decreased vimentin expression and prevented cells at the leading edge from extending vimentin-rich lamellipodia, resulting in impaired wound closure [60]. In a more recent study, Berr et al. showed that spheroids constructed from cells derived from Vim-/- genetically engineered mouse model have significantly lower growth than Vim+/+ spheroids in collagen I [61]. Similarly, Sandrine et al. showed that IF-depletion glioblastoma spheroids have strongly decreased cell invasion in 75% Matrigel [62]. Our result here confirmed the established phenotype of Vim+/+ cells: they exhibit enhanced migratory behavior collectively.

While our results are consistent with the picture that vimentin enhances collective cell migration, there are several aspects of our findings that could be considered surprising for at least three reasons based on cell studies. First, recent work has shown vimentin diminishes actin stress fiber contractility by deactivation of RhoA through its guanine nucleotide exchange factor GEF- H1 [24]. This study demonstrated loss of vimentin does not necessarily diminish cell contractility and may even promote it. We do note that recent work shows that vimentin’s effects on 3D cell contractility are matrix stiffness dependent, such that this effect is diminished on soft gels [25]. Second, vimentin has been shown to decrease cell speeds in confining 3D spaces. For individual cells in micro-channels and collagen network, vimentin diminishes the speed of individual cells moving in micro-channels and collagen networks, which is opposite of what we find for the effects of vimentin on the collective fibroblast aggregates here. For individual cells, the decrease in cell speed has been attributed to resistance of deforming the vimentin network to squeeze through small spaces, which confers protection to the cell nucleus [22]. Reason three is connected to the first two. We find here that vimentin increases nuclear elongation, whereas prior studies indicate that vimentin reduces nuclear deformations and elongation, albeit under conditions of extreme external confinement as single cells squeezing through 3 μm collagen pores or cells are subject to uniaxial pressure [63].

To interpret our result, we suggest a minimal model of collective cell migration in fibrous environments that captures the effects of vimentin. In this picture, vimentin has at least two different yet complementary effects, namely roles in collagen degradation and enhanced cell contractility via increased polarization.

First, as shown in figure 5A, collective fibroblast migration through collagen networks is MMP-dependent, at least for the collagen concentration used here where the pore size is 2-5 µm, small compared to the size of the cell. The role of vimentin in collagen degradation has recently been reported. Using wild-type and vimentin-null mEFs, Ostrowska-Podhorodecka et al showed that vimentin’s interaction with myosin 10 was required for pericellular collagen degradation by MT1-MMP. In particular, vimentin deletion reduced MT1-MPP transport toward cellular extensions, down-regulated collagenolytic activity, reduced collagen fiber alignment and cell invasion of the matrix [26]. Further, these effects required physical vimentin networks not just soluble vimentin units, as shown using the ULF cells, consistent with the reduced migration of ULF cells found here (Fig. 4). This effect could explain the reduced cell invasion seen in the spheroids of mixed vimentin-null – MDA-MB-231 cells in which some of the proteolytic-capable MDA-MB-231 are replaced with proteolytic-deficient vimentin-null cells. Further, as shown in Fig. 5D, vimentin-null cells from the spheroids are capable of moving out into the matrix, once they can reach large enough channels, generated here by laser ablation.

A second and unexpected finding is the evidence that vimentin increases fluidization of the collective fibroblast spheroids, as shown by the increase in nuclear elongation in Fig. 6B. Vimentin is known to increase cell polarization through its interactions with microtubules and actomyosin networks [64–69]. Interestingly, we have found when cells are geometrically constrained to polarized shapes on micropatterned substrates, vimentin increases the transmission of internal acto-myosin based forces to the cell nucleus, leading to larger deformations and elongations of the nucleus [70]. One possibility for vimentin’s differing effects on nuclear shape is that there are differences in how vimentin mediates passive outside-in applied forces versus active internally-generated forces. Namely, vimentin may resist large external deformations of the cell from outside boundaries that decrease nuclear deformations, it may also assist in the transmission of internal actomyosin contractility that deforms the nucleus by interlinking to the actomyosin network and to the nuclear envelope through the LINC complex via nesprin-3. Another factor influencing nuclear shape is cell shape, such as those driven by differences in cell-cell adhesion; however, our cell-sorting experiments do not support a significant differential adhesion between the cells (SI Fig. 5).

The impact of vimentin on nuclear shape has interesting implications for the mechanical state of spheroids. Recent work by Grosser et al. showed that elongated nuclear shapes correlate with elongated cell shapes and are a marker of the degree of tissue fluidity and cell mobility within tissue [41]. With respect to invasive cancers, cancerous samples or aggregates of cancer cells are more fluidized compared to healthy solid-like cell aggregates [41, 71], suggesting a change in mechanical state that correlates with metastatic potential. Further, a recent computational model based on a 3D vertex cell model embedded in a collagen spring network has been proposed, which predicts that a more fluidized tumor spheroid will pull on and densify collagen fibers more closely around a spheroid, increasing the spheroid contractility, consistent with enhanced migration into the network[49]. Taken together, the data in Fig. 7 is consistent with the correlation between increased vimentin expression and cancer progression and suggests a mechanism by which vimentin promotes metastatic spread of cancer through modifying nuclear shape and tissue fluidity.

Altogether, our experimental results motivate a physical mechanism in which vimentin-mediated collagen degradation is a pre-requisite for collective fibroblast migration in collagen networks and loss of vimentin-mediated collagen degradation is a limiting factor. In this sense, our results suggest vimentin’s role in MMP-transport and collagen degradation is a predominate effect in inhibiting collective migration in ECM networks. However, a secondary effect of vimentin is the polarization of cells associated with the fluidization of the spheroids, which promotes collective cell expansion and may aid in collective spheroid contractility. One area of future exploration is to address the relative extent of these two effects on collective migration and if, for instance, collagen-degradation was rescued in vimentin-null cells and ULF cells, whether or not their collective migration through collagen would phenocopy the wild-type expansion.

It is not surprising that collagen degradation is necessary to generate the space needed for collective cell migration though collagen and many studies have highlighted the role of collagen degradation in spheroid expansion [33, 35, 72, 73], but the primary role of vimentin in collagen degradation may be surprising, given the focus of vimentin’s role in cytoskeletal mechanics. Our studies highlight the importance of vimentin’s dual role in mechanical and non-mechanical signaling functions in recognizing its full impact and multifactored functions in cells and tissues. Further, the differing effect of vimentin on single cell versus collective cell migration highlights the influence of cell-cell contacts and proteolytic activity during collective migration through fibrous networks.

In summary, we identify novel effects of vimentin on collective cell migration. Collectively, vimentin impacts fibroblast spheroid matrix remodeling and spheroid fluidity, relevant to collective cell migration through soft connective tissue. Future studies will be needed to determine the specific molecular mechanisms underlying vimentin’s role in collective migration and to decipher its influence at multiple different length and times scales associated with migration, ranging from ECM remodeling, cell shape, and nuclear positioning.

Fibroblasts are known for their ability to remodel tissue and to increase the motion of cancer cells through various factors, including contraction of the ECM network and degradation of the ECM through MMPs [74]. Our results thus underscore that vimentin networks are essential for the critical functions of fibroblasts in health and disease, which originates from vimentin’s signaling and mechanical roles. Understanding the mechanisms by which cells collectively navigate and remodel their matrix environments will help identify how migration is altered during disease and strategies to mitigate it.

## Materials and Methods

### Cell Culture

Vimentin Wildtype (Vim+/+) and vimentin knockout (Vim-/-) mouse embryonic fibroblasts (MEFs) were gifted by J. Eriksson from Abo Akademi. Vimentin unit length filament (VimULF) cells were gifted by Goldman lab from Northwestern University. MDAMB231 human breast cancer cells were given by Mingming Wu lab from Cornell University. All cells were cultured in Dulbecco’s Modified Eagle’s Medium + 4.5g/L glucose + 2nM L-glutamine + sodium pyruvate (DMEM, Gibco) and supplemented with 10% fetal bovine serum (FBS, Hyclone), 1% non-essential amino acid (Fisher Scientific), 1% penicillin/streptomycin (Fisher Scientific) and 25 nM of HEPES (Fisher Scientific). Cell cultures were maintained at 37 ^0^C and 5% CO2. Cells were passaged at confluency 70-80% and medium was replenished every 3 days.

### Spheroid Culture

To create spheroid invasion assay, first we made microwells by adapting PRIMO (Alvéole) spheroid protocol [75]. In brief, 1% agar spin-coated glass slide was glued to the bottom of a 35mm petri dish to create a non-adherent surface. Next PDMS stencils were assembled on top of the coated glass surface to create a hydrogel PDMS chamber. Then 50% 4-Arm-PEG-Acryl in PLPP Classic (PRIMO) solution was injected into the PDMS chamber. Next, the solution was exposed to UV light with the PRIMO device using Leonardo software with the appropriate photomask (well radius = 200µm, well height = 250µm, 45mJ/mm^2^ for 20 seconds). After the polymerization, the stencils were then removed and the microwells were washed under PBS 3x, exposed under UV light for 15 minutes and then incubated in cell media for 20 minutes. Finally, cells were plated at high density 80000 cells/dish in DMEM/F12 (10% FBS, 1x Pen/Strep) to ensure the cells would get into the microwells instead of rolling off to the empty area. Cell aggregates were incubated at 37°C with 5% CO_2_ for 2 days before harvesting.

To perform spheroid spreading assays, cell aggregates were first generated using the procedure described by Ibidi® [76]. Briefly, 96-well flat-bottom plates (Corning) were coated with 40 µl of 1% agarose solution diluted with PBS (Fisher Scientific). Mouse embryonic fibroblasts (Wild-type and vimentin-null) were then plated at density of 30000 cells/well in 100 µl of DMEM/F12 (10% FBS, 1xPen/Strep). Cell aggregates were incubated at 37°C with 5% CO_2_ for 2 days before harvesting.

For co-culture spheroids, cells were first live stained with CellTracker^TM^ (ThermoFisher) before transfer to microwells to form spheroids.

### Collagen Gel

Collagen gels were prepared using a slightly modified protocol from Ibidi® [77]. Collagen type I, rat tail 5mg/ml (Ibidi) were diluted on ice to 1.5mg/ml with 6.67% of 5x DMEM (Fisher scientific), 10% FBS in 1x DMEM. The pH was adjusted to 7.4 using 1M NaOH and sodium bicarbonate. Cell aggregates were added to the collagen solution before it solidified. The collagen solution is then pipetted on 1% agarose coated 24-well flat bottom plates and let polymerized for 40 minutes in incubator at 37oC, 5% CO2 and 100% humidity. DMEM complete media was added on top of the fully polymerized collagen gel.

### Collagen Gel Rheology Characterization

All rheology tests were performed with a Malvern Panalytical Kinexus Ultra+ rheometer (Malvern Panalytical) using a 20 mm parallel plate geometry. Collagen hydrogel solutions were mixed and immediately cast onto the bottom plate of the rheometer. The upper plate was then lowered to a gap height of 1mm. The gelation was tracked over the course of 30 min using a single frequency oscillatory sweep (f = 1Hz, ε = 2%) test at a temperature of 37°C. The gelation was monitored by the G’ and G’’, which reached a plateau value at around 20 min. For compression tests, samples were subjected to stepwise lowering of the top plates, corresponding to increasing steps of 4% compressive strain up to 20% strain. At each compressive strain %, the sample was allowed to equilibrate for 5min. All compression tests were performed with a continuous single frequency oscillatory sweep (f = 1Hz, ε = 2%). These compression tests were also used to estimate the Young’s Modulus E of the sample. First, the final normal force value at each compressive step was used to calculate axial stress. Then plotting axial stress vs axial strain, a linear fit was applied where the slope provides an estimate of Young’s Modulus E.

### Collagen Gel Network Characterization

For analysis of the collagen gel network, reconstituted collagen gels were also prepared using the recipe above without spheroids. Collagen network was imaged with confocal reflection microscopy (CRM) on the LSM 980 (Zeiss) in reflected mode with 40x water immersion objective (LD LCI Plan-Apochromat 40x/1.2 Imm Corr DIC M27, Zeiss). The excitation laser wavelength is 488nm and backscatter was collected via beam splitter BC80/20. Confocal image stacks (50 µm) of four samples were captured at z-resolution of 0.25 µm.

Pore size of the collagen gel was analyzed by a slight modification of the FIbeR Extraction MATLAB code used in Stein et al. [78] to include bubble analysis algorithm by Munster & Fabry et al. [79]. In brief, the 3D image stack is first smoothed with a gaussian filter then binarized with appropriate threshold. Next, minimal distance from a fiber pixel to a background pixel is calculated using MATLAB’s bwdist.m function in the image processing toolbox. The distance is then smoothed with a Gaussian filter. From this point, we feed the smoothed distance function to bubble analysis code and extract the pore size.

Collagen network topography was also analyzed using topography-analysis code by Katja et. al. [80]. Here, we again extract the pore size, fiber diameter, fiber orientation, fiber density and network anisotropy.

Collagen compaction was found with method previously used in [81]. In brief, the mean intensity of the confocal reflectance image of collagen fibrils in fixed area around the core spheroid was computed and then normalized against the mean intensity of the collagen fibrils at the corners (far away from the spheroid). (Fig S4).

For measuring collagen alignment index, we utilized a method that has been done previously in [82] with some modification due to strong scatter signal from the core of the spheroid. Nine sub regions surrounding the core spheroid of the confocal reflectance image of the collagen fibrils are cropped and transformed with 2D Fast Fourier Transform (MATLAB). The transformed image was then replotted as a function of radial sum of the intensity as a function of angle. The alignment index was defined based on higher pixel intensities in a specific angle, which was related to the orientation of collagen fibers in the corresponding direction. Quantification was done by calculation of the area under the intensity curve within ± 10 degrees of the peak using a self-written MATLAB script. (Fig S4)

### Immunofluorescence Staining

Spheroids embedded in collagen were washed 3x in PBS supplemented with CaCl_2_ and MgCl_2_ and fixed with 4% paraformaldehyde (Sigma) for 30 min at room temperature, permeabilized with 0.5% Triton X-100 in PBS (PBS-T) at 4^0^C for overnight. Next, spheroids were stained with primary antibody (anti-rabbit Vimentin) in blocking buffer at 4^0^C overnight. After that, samples were washed 5 times (10 minutes each) and then incubated in optical clearing solution (80% Glycerol in PBS) at 4^0^C overnight [83]. Next, spheroids were stained with secondary antibody (anti-rabbit 488) in blocking buffer at 4^0^C overnight. Blocking buffer includes 1% Tween-20, 1% Triton-X, 1% DMSO, 0.1% Sodium Azide and 1% BSA in PBS solution (adapted from Adhya et al. [84]). Samples were then again washed 5 times and incubated in optical clearing solution at 4^0^C overnight. Next, spheroids were counterstained with Rhodamine Phalloidin (Alexa) at 1:300 dilution and DAPI 2 μg/ml in 40% DMSO:PBS solution at 4^0^C overnight. Finally, samples were washed 5 times and incubated in optical clearing solution at 4^0^C overnight before getting mounted for imaging.

### Morphological quantification

After 3 hours inside the incubator, spheroids in collagen gel are imaged for 48 hours with phase-contrast widefield using a Plan Fluor 10× objective on Nikon Eclipse Ti inverted microscope equipped with an Andor Technologies iXon em+ EMCCD camera (Andor Technologies). Aggregates were maintained at 37 °C and 5% CO2 using a Tokai Hit (Tokai-Hit) stage top incubator. Cross-sectional area of the spheroid (core and connecting strands) was preprocessed with custom MATLAB code and traced using ImageJ [85].

### Nucleus Segmentation

Segmentation was done using the segmentation pipeline XPIWIT [86] which is a XML pipeline wrapper for ITK segmentation software [87]. The software package is available at https://bitbucket.org/jstegmaier/xpiwit/downloads/ and it has been verified to work very well on nuclear segmentation [88]. Raw DAPI channel of the spheroid confocal images were obtained and used as an image input for the software. The XML pipeline chosen was the default TWANGSegmentation provided. Custom MATLAB script was used to generate input file and PowerShell script to batch run XPIWIT for each confocal image stack. Output of XPIWIT was then queries into Microsoft Excel Data Model (Data Analysis Plugin) for data cleanup: removing duplicates, filter out nucleus volume that are too small (image artifact). Nucleus aspect ratio is calculated from extracted major and minor axis and plotted with Jupyter Notebook (Python). Statistic comparison between population was done using MATLAB.

### Traction Force Microscopy

FluoroSphere (2 µm) stock solution [(ThermoFisher) was added to the collagen gel mixture (1.5mg/ml) at 1:50 ratio. Spheroids were prepared as previously described and transferred into collagen gel mixture with fluorescent beads. Collagen gel was then polymerized in the incubator for 1 hour and before cell media was added on top. After 3 hours, spheroid was then mounted and imaged every 15 minutes for 48 hours using Plan Fluor 10× objective on Nikon Eclipse Ti inverted microscope equipped with an Andor Technologies iXon em+ EMCCD camera (Andor Technologies) and Lambda LS (Sutter Instrumentation Company) for fluorescent excitation. Sample were maintained at 37 °C and 5% CO2 using a Tokai Hit (Tokai-Hit) stage top incubator. Traction force was calculated using jointforces package provided by Christoph et al. at https://github.com/christophmark/jointforces. In brief, beads displacement was computed using OpenPIV [89] provided via jointforces package. 1.5 mg/ml collagen lookup table was created using jointforce material simulation with parameters found from collagen rheology characterization. Contractility was then finally reconstructed using PIV result output and material simulation lookup table.

### Laser-ablated channels

Laser ablation of microchannels in cell encapsulated collagen ECM was performed using a femtosecond (fs) laser platform, which combined a Ti:Sapphire fs laser (Coherent, Chameleon, USA) with a Zeiss Microscope (Observer Z1, Germany). In this setup, 800 nm fs-laser beam with a repetition rate of 80 MHz was focused into the crosslinked collagen using a water immersion objective (40, NA=0.8, Leica). Visual basic code was employed to control the movement of microscope stage and the shutter, allowing for controlled scanning of laser focus within the collagen to ablate 2D microchannels. The laser dosage needed for ablation of collagen concentration was determined by varying laser power and scanning speed of the laser focus. Specifically, a laser power of 1.2 W, measured just before the objective, and scanning speed of 50 μm/sec was used to ablated microchannels.

### Statistics

Data is presented as mean ± standard error of the mean (SEM). Statistical analyses were performed using GraphPad Prism 9 (GraphPad Software, Inc.) unless stated. Comparison between Vim+/+, Vim-/-and VimULF are done using Welch ANOVA test (multiple comparison). Denotations: **** p<0.0001, *** p<0.001, ** p<0.01, * p<0.05. n denotes number of experiments and N denotes sample population of a phenotype.

## Acknowledgements.

We thank Profs. Paul Janmey, Ben Fabry, Vivek Shenoy, and Lisa Manning for useful discussions on this project. Support for this project was provided by an NIH grant R35 GM142963 awarded to AP. This project was also supported by NSF grant NSF-PoLS-2014192. TZ acknowledges financial support from the NSFC/China via award 22303051. We are thankful to the National Institutes of Health (R21 GM141573-01) for the financial support for this project.

